# The Arabidopsis *katamari2* Mutant Exhibits a Hypersensitive Seedling Arrest Response at the Phase Transition from Heterotrophic to Autotrophic Growth

**DOI:** 10.1101/2023.11.05.565727

**Authors:** Chika Hosokawa, Hiroki Yagi, Shoji Segami, Atsushi J. Nagano, Yasuko Koumoto, Kentaro Tamura, Yoshito Oka, Tomonao Matsushita, Tomoo Shimada

## Abstract

Young seedlings use nutrients stored in the seeds to grow and acquire photosynthetic potential. This process, called seedling establishment, involves a developmental phase transition from heterotrophic to autotrophic growth. Some membrane-trafficking mutants of *Arabidopsis thaliana* (Arabidopsis), such as the *katamari2* (*kam2*) mutant, exhibit growth arrest during seedling development, with a portion of individuals failing to develop true leaves on sucrose-free solid medium. However, the reason for this seedling arrest is unclear. In this study, we show that seedling arrest is a temporal growth arrest response that occurs not only in *kam2* but also in wild-type Arabidopsis; however, the threshold for this response is lower in *kam2* than in the wild type. A subset of the arrested *kam2* seedlings resumed growth after transfer to fresh sucrose-free medium. Growth arrest in *kam2* on sucrose-free medium was restored by increasing the gel concentration of the medium or covering the surface of the medium with a perforated plastic sheet. Wild-type Arabidopsis seedlings were also arrested when the gel concentration of sucrose-free medium was reduced. RNA sequencing revealed that transcriptomic changes associated with the rate of seedling establishment were observed as early as 4 days after sowing. Our results suggest that the growth arrest of both *kam2* and wild-type seedlings is an adaptive stress response and is not simply caused by the lack of a carbon source in the medium. This study provides a new perspective on an environmental stress response under unfavorable conditions during the phase transition from heterotrophic to autotrophic growth in Arabidopsis.

## Introduction

The life of a seed plant can be divided into several developmental stages: germination, vegetative growth, flowering, and senescence. After germination, the seedling uses the nutrients stored in the seeds to grow and acquire photosynthetic capacity. This process, called seedling establishment, involves a developmental phase transition from heterotrophic to autotrophic growth (Ha et al., 2017). *Arabidopsis thaliana* (Arabidopsis) seeds store enough nutrients to allow the seedlings to grow for 4–5 days (Kircher and Schopfer, 2012). The transition of the growth phase is crucial for plant survival and is regulated by environmental factors, such as light, temperature, minerals, and water availability, as well as internal factors, such as storage substances and phytohormones (e.g., auxin and abscisic acid) (Gommers and Monte, 2018; Vidal et al., 2014). For example, germination in the dark results in etiolated seedlings that cannot transition to autotrophic growth, and metabolic deficiencies of storage substances also cause seedling arrest.

Arabidopsis seeds accumulate lipids and proteins as the major storage materials, which are metabolized during heterotrophic growth after germination and used as carbon and nitrogen sources (Baud et al., 2008). Storage lipids accumulate as triacylglycerols in organelles called lipid droplets (oil bodies). Triacylglycerols are degraded to fatty acids by the lipolytic enzyme SUGAR-DEPENDENT1 (SDP1) during germination (Kelly et al., 2011). The fatty acids are further metabolized in peroxisomes and mitochondria, and used as a substrate for glycogenesis. Storage proteins, which accumulate in protein storage vacuoles, are biosynthesized as precursors in the rough endoplasmic reticulum; transported by membrane trafficking via endomembrane systems such as the Golgi apparatus, *trans*-Golgi network, and pre-vacuolar compartment to reach the vacuole; and converted to their mature form (Shimada et al., 2018). Storage proteins are degraded during germination and mainly used as a nitrogen source for seedling growth (Kaur et al., 2021). In Arabidopsis, storage proteins are also important carbon sources that can be incorporated into the glycogenesis pathway (Eastmond et al., 2015).

Membrane trafficking is responsible for the intracellular transport of proteins and other macromolecules and plays important roles in plant growth, development, and stress responses (Surpin and Raikhel, 2004). Membrane vesicular trafficking, an evolutionarily conserved process in eukaryotes, involves four steps: (1) budding of coat protein-coated transport vesicles from the donor membrane, (2) intracellular transport of transport vesicles from the donor organelle to the target organelle, (3) tethering of transport vesicles to the target organelle membrane via tethering factors, and (4) membrane fusion of transport vesicles and target organelles via soluble NSF attachment protein receptors (SNAREs). PIN-FORMED (PIN) family proteins are membrane-trafficking cargo proteins with important roles in plant growth and development (Adamowski and Friml, 2015). PIN proteins transport the phytohormone auxin and are unevenly distributed in specific regions of the plasma membrane to form auxin concentration gradients in the plant body. Membrane trafficking plays important roles in various auxin-mediated morphogenetic and environmental responses, such as embryogenesis, vascular development, organogenesis, bending, apical bud dominance, and shade avoidance responses.

We previously described Arabidopsis KATAMARI2 (KAM2), a membrane-trafficking factor involved in endosome formation and the intracellular trafficking of vacuolar proteins (Tamura et al., 2007). *kam2* mutants have mutations in the same gene (At2g26890) as *gravitropism defective 2* (*grv2*) mutants, and some *grv2* individuals exhibit arrested seedling growth on sucrose-free solid medium (Silady et al., 2008). Similar to *kam2*/*grv2*, several other membrane-trafficking mutants exhibit sucrose-dependent seedling establishment, including *protein affected trafficking4* (*pat4*) (Zwiewka et al., 2011), *sorting nexin1* (*snx1*) (Kleine-Vehn et al., 2008), and *maigo1*/*vacuolar protein sorting29* (*mag1*/*vps29*) (Thazar-Poulot et al., 2015). However, the physiological significance of this seedling arrest has not been explored. Here, we investigated features of this growth arrest phenotype using the *kam2* mutant as a model. Our results suggest that seedling arrest is not caused by the lack of a carbon source; instead, it appears to be an environmental stress response that also occurs in wild-type Arabidopsis.

## Results

### *kam2* seedlings show growth arrest on sucrose-free medium but resume growth when transplanted to fresh sucrose-free medium

We investigated whether the transfer-DNA (T-DNA) insertion alleles of *kam2* exhibit a seedling arrest phenotype similar to that observed in the allelic mutant *grv2* (Silady et al., 2008). On sucrose-containing solid medium (0.5% gellan gum), almost all *kam2-2* and *kam2-4* T-DNA insertion mutants exhibited normal growth (Fig. 1A, B). By contrast, when *kam2-2* and *kam2-4* mutants were grown on sucrose-free solid medium (0.5% gellan gum), most individuals stopped growing after germination, with short roots and no true leaf development (Fig. 1A). By 2 weeks after germination, only 25% and 34% of the *kam2-2* and *kam2-4* seedlings had developed true leaves and elongated roots, respectively (Fig. 1B). The growth arrest observed for *kam2* under these conditions is generally consistent with that reported previously for *grv2* (Silady et al., 2008). The medium was solidified with gellan gum in our experiment, but with agar in the previous report. Thus, the growth arrest phenotype of *kam2*/*grv2* seedlings is independent of the gelling agent used in the medium. The percentage of seedling establishment varied somewhat depending on the production lot of the medium and lots of seeds (different storage periods and/or conditions).

**Fig. 1.**
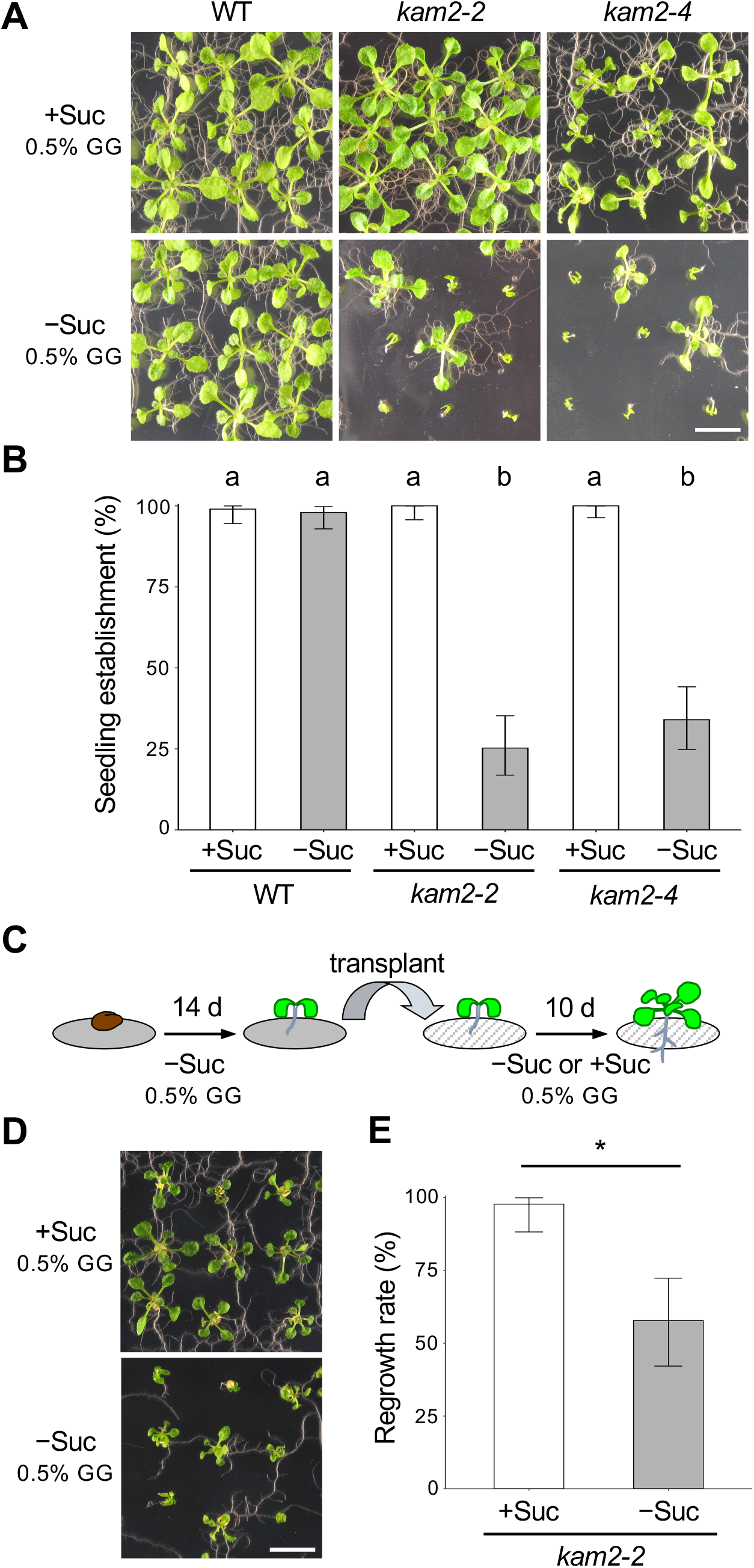
The *kam2* mutants exhibit defects in seedling establishment. (A) Images of 14-day-old wild-type (WT), *kam2-2*, and *kam2-4* seedlings grown on solid culture medium (0.5% gellan gum, GG) supplemented with (+Suc) or without (–Suc) 1% sucrose. Bar, 1 cm. (B) Quantitative analysis of seedling establishment in 14-day-old WT, *kam2-2*, and *kam2-4* seedlings as shown in (A). Error bars show binomial 95% confidence intervals. Different letters indicate significant differences between the six samples at *p* < 0.01 (Fisher’s exact test with Bonferroni multiple testing correction). *n* = 100 (WT, +Suc), 99 (WT, –Suc), 84 (*kam2-2*, +Suc), 95 (*kam2-2*, –Suc), 99 (*kam2-4*, +Suc), and 100 (*kam2-4*, –Suc). (C) Schematic representation of the transplantation experiment. Fourteen-day-old seedlings grown on solid culture medium without sucrose were transferred to fresh solid medium with or without 1% sucrose and grown for an additional 10 days. (D) Images of 24-day-old *kam2-2* seedlings transferred to culture medium (0.5% GG) with (+Suc) or without (–Suc) 1% sucrose. Bar, 1 cm. (E) Quantitative analysis of regrowth rate in transferred *kam2-2* seedlings as shown in (D). Error bars show binomial 95% confidence intervals. Asterisk indicates a significant difference (Fisher’s exact test *p* < 0.01). *n* = 45 (+Suc), 45 (–Suc).

Next, we transplanted arrested seedlings onto sucrose-containing medium to test their growth. In a previous study, *grv2* seedlings were transplanted within 6 days after germination, which may have been too early to observe seedling arrest (Silady et al., 2008); here, we transplanted the seedlings at 2 weeks after germination, when arrest was clearly visible (Fig. 1C). We also transplanted the arrested seedlings onto sucrose-free medium as a control. When *kam2-2* seedlings that had been grown for 14 days on sucrose-free medium (0.5% gellan gum) and had not developed true leaves were transferred to sucrose-containing solid medium (0.5% gellan gum), seedling growth resumed, and almost all seedlings had developed true leaves and elongated roots by 10 days after transfer (Fig. 1D, E). Surprisingly, even when we transferred *kam2-2* seedlings to sucrose-free solid medium (0.5% gellan gum), roughly half of the seedlings resumed growth with elongated roots and developed true leaves (Fig. 1D, E). Small environmental changes during the transfer may have triggered this resumption of growth. These results indicate that the seedling arrest of *kam2* on sucrose-free solid medium may be triggered by environmental factors in addition to a genetic factor.

### The gel concentration of the medium affects the establishment of wild-type and *kam2* seedlings

To investigate the cause of seedling arrest in *kam2*, we varied the conditions of the medium. Since it was previously reported that varying the gel hardness of the solid medium changed the growth of plants (Fukuda et al., 2016), the gellan gum concentration of sucrose-free solid medium was changed. In addition to the routinely used solid medium (0.5% gellan gum), we grew wild-type and *kam2* plants on 0.2% and 1% gellan gum sucrose-free media (Fig. 2A). The seedling establishment rates of the wild type were 99% on 1% gellan gum medium and 93% on 0.5% gellan gum medium but decreased to 45% on 0.2% gellan gum medium (Fig. 2B). By contrast, the seedling establishment rates of *kam2-2* were 41% on 0.5% gellan gum medium but decreased to 29% on 0.2% gellan gum medium and increased to 82% on 1% gellan gum medium (Fig. 2B). These results suggest that the seedling arrest can occur not only in the *kam2* mutant but also in wild-type Arabidopsis plants on softer media and that the *kam2* mutant is more sensitive to the growth substrate than the wild type.

**Fig. 2.**
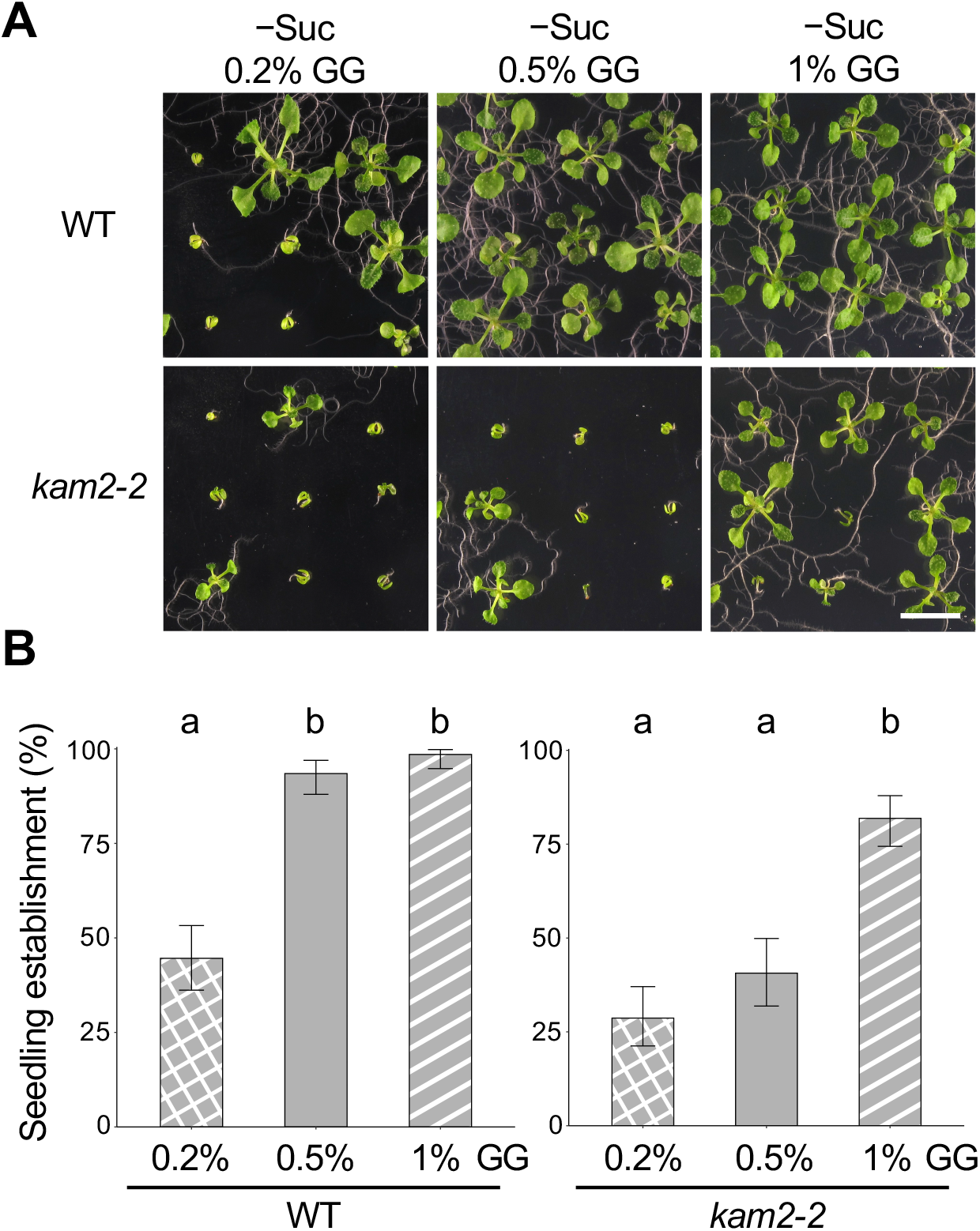
The gel concentration in solid medium affects seedling establishment in wild-type and *kam2* seedlings. (A) Images of 14-day-old wild-type (WT) and *kam2-2* seedlings grown on solid culture medium containing 0.2, 0.5, or 1% gellan gum (GG) without sucrose (–Suc). Bar, 1 cm. (B) Quantitative analysis of seedling establishment in 14-day-old WT and *kam2-2* seedlings as shown in (A). Error bars show binomial 95% confidence intervals. Different letters indicate significant differences between three samples within the same genotypes at *p* < 0.01 (Fisher’s exact test with a Bonferroni multiple testing correction for each genotype). *n* = 139 (WT, 0.2%), 138 (WT, 0.5%), 136 (WT, 1%), 136 (*kam2-2*, 0.2%), 123 (*kam2-2*, 0.5%), and 138 (*kam2-2*, 1%).

### Placing a plastic sheet on solid medium partially rescues the growth arrest of *kam2*

A previous study showed that using growth media with low gellan gum concentrations tends to bring the shoots in contact with the medium (Fukuda et al., 2016), suggesting that the physical contact of the shoots with the media may be responsible for the growth arrest phenotype. To investigate the possibility that physical contact between the shoots and the medium is the cause of seedling arrest, we conducted an experiment using perforated plastic sheets to cover the medium. We placed plastic sheets with 1-mm-diameter holes on top of solid medium and sowed the seeds in the holes so that the aboveground parts of the seedling were less likely to come in contact with the medium (Fig. 3A). The seedling establishment rates of the *kam2* mutant on sucrose-containing and sucrose-free medium without the plastic sheet (0.5% gellan gum) were 100% and 26%, respectively (Fig. 3B). When we placed a plastic sheet on top of solid, sucrose-free medium (0.5% gellan gum), the proportion of *kam2* individuals growing normally increased to 81% (Fig. 3B). However, established seedlings of both wild-type and *kam2* plants grown on plastic sheets were slightly smaller than those of seedlings grown on the control medium without plastic sheets (Fig. 3A). The wild type showed more than 95% seedling establishment under all conditions (Fig. 3B). These results strongly suggest that the major cause of seedling arrest in the *kam2* mutant is contact between the shoots and the medium.

**Fig. 3.**
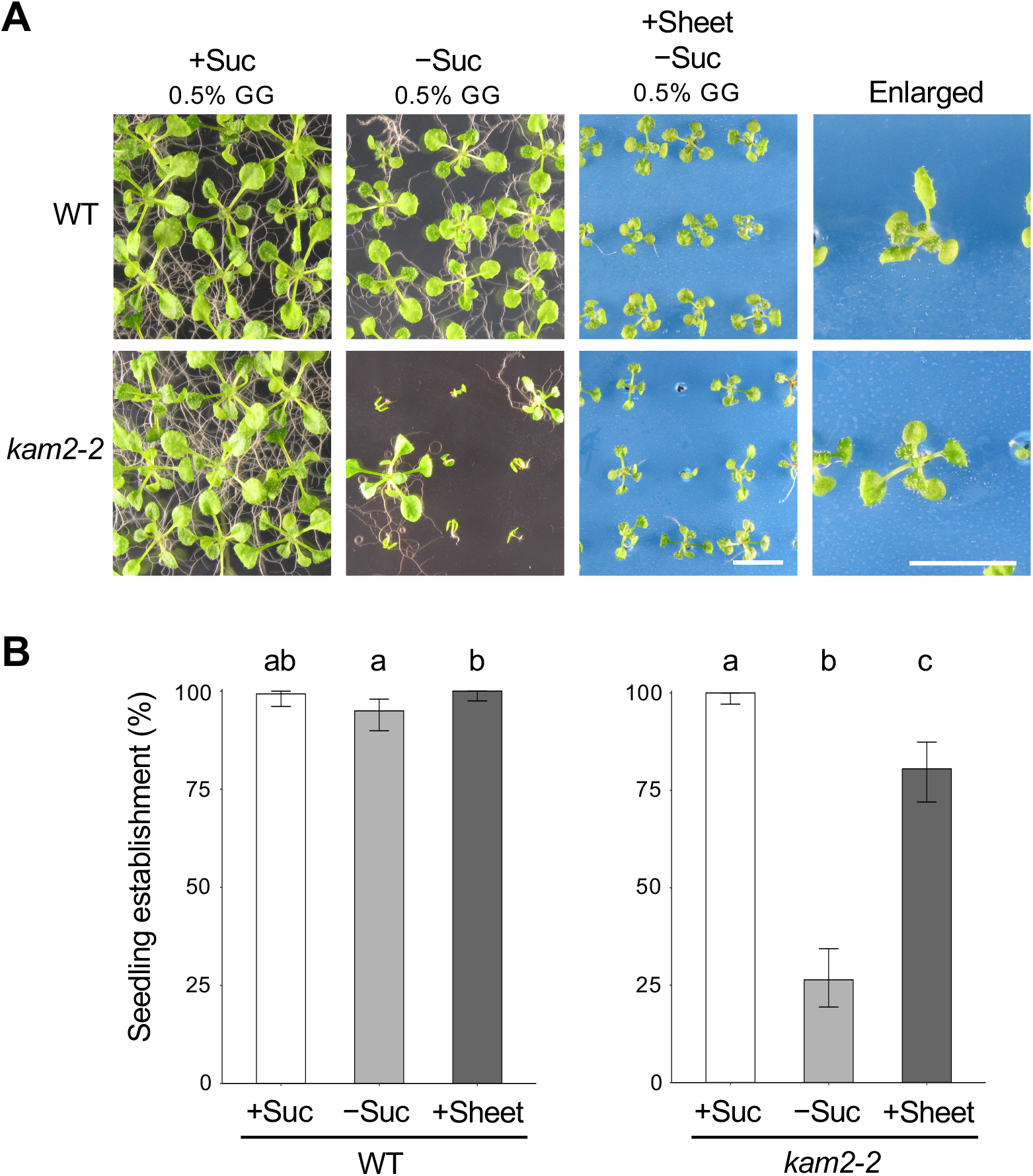
Applying a plastic sheet on solid medium rescues the seedling establishment defect in *kam2*. (A) Images of 14-day-old wild-type (WT) and *kam2-2* seedlings grown on solid culture medium (0.5% gellan gum, GG) lacking sucrose (–Suc) and covered with (+Sheet) or without a blue plastic sheet. The rightmost panels are enlarged images of seedlings on plastic sheets. The leftmost panels are control seedlings grown on medium (0.5% GG) containing 1% sucrose (+Suc). Bars, 1 cm. (B) Quantitative analysis of seedling establishment in 14-day-old WT and *kam2-2* seedlings as shown in (A). Error bars show binomial 95% confidence intervals. Different letters indicate significant differences between three samples within the same genotypes at *p* < 0.05 (Fisher’s exact test with a Bonferroni multiple testing correction for each genotype). *n* = 140 (WT, +Suc), 139 (WT, –Suc), 146 (WT, +Sheet), 127 (*kam2-2*, +Suc), 144 (*kam2-2*, –Suc), and 113 (*kam2-2*, +Sheet).

### Growth arrest of wild-type seedlings on softer medium is restored by the addition of sucrose or by placing a plastic sheet on the solid medium

The above experiments revealed that seedling arrest, which was initially thought to be a phenotype of the *kam2* mutant, was also induced in wild-type Arabidopsis plants grown on a softer solid medium (0.2% gellan gum). The growth arrest induced in *kam2* seedlings on normal solid medium (0.5% gellan gum) could be alleviated by adding 1% sucrose to the medium, increasing the concentration of gellan gum in the medium (1% gellan gum), or placing a perforated plastic sheet on the medium. Therefore, we examined whether the growth arrest of wild-type seedlings on medium containing 0.2% gellan gum could also be alleviated under these treatments (Fig. 4A, B). The seedling establishment rates of the wild type and *kam2-2* mutant were 66% and 34%, respectively, on 0.2% gellan gum sucrose-free medium and increased to 86% and 52%, respectively, on the same medium covered with perforated plastic sheets (Fig. 4B). Moreover, the seedling establishment rates of both the wild type and *kam2-2* mutant were 100% on 0.2% gellan gum medium supplemented with 1% sucrose (Fig. 4B). These results indicate that the growth arrest of wild-type Arabidopsis on a softer medium can be alleviated by medium conditions, as can the growth arrest of *kam2* mutants. Thus, the cause of growth arrest in *kam2* mutants and the wild type appears to be the same, although the threshold is different.

**Fig. 4.**
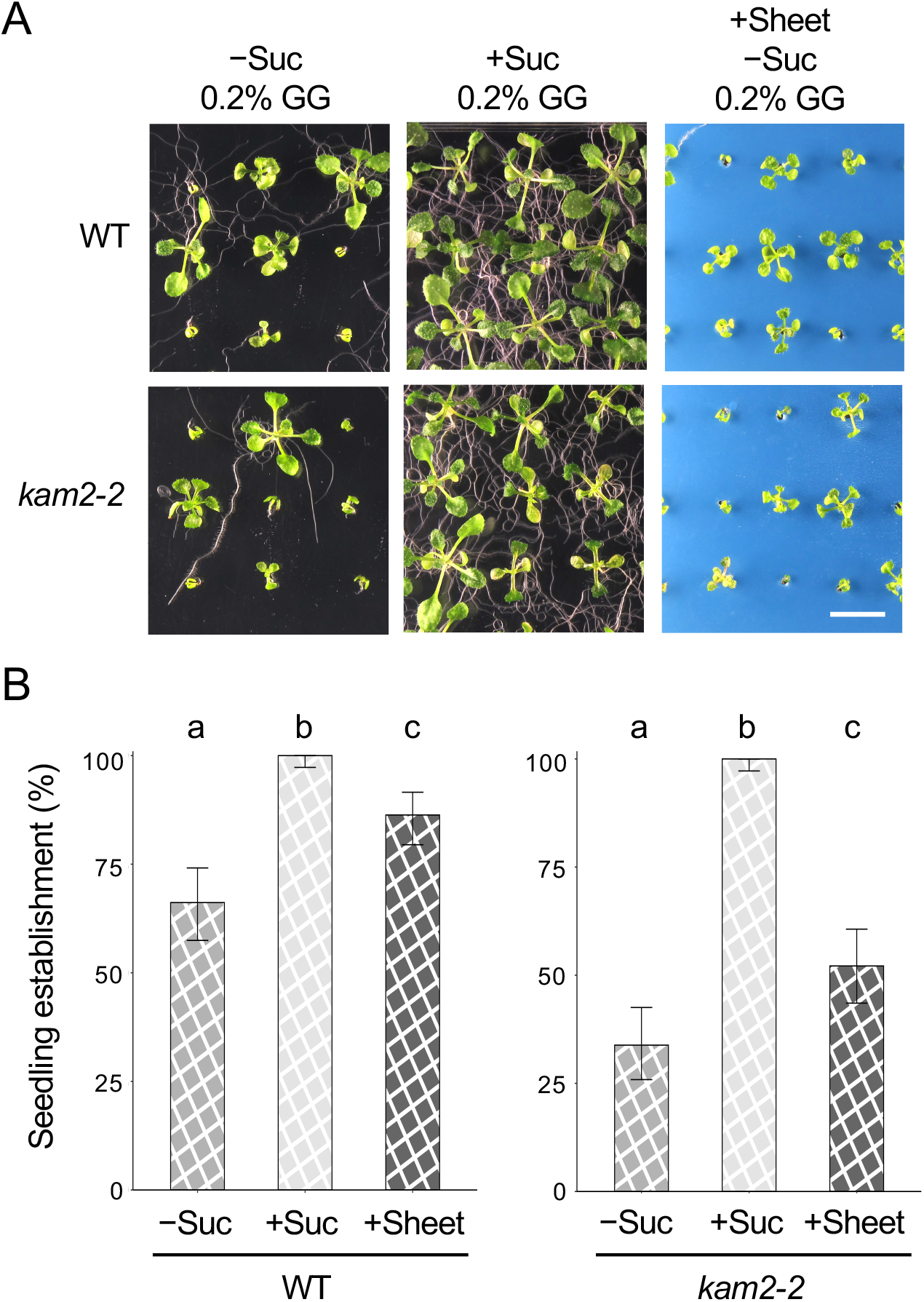
Seedling establishment is restored in wild-type Arabidopsis plants growing on soft solid medium if sucrose is added to the medium or a plastic sheet is placed on the surface. (A) Images of 14-day-old wild-type (WT) and *kam2-2* seedlings grown on solid culture medium (0.2% gellan gum, GG) with (+Suc) or without (–Suc) 1% sucrose, or without 1% sucrose and covered with a blue plastic sheet (+Sheet). Bar, 1 cm. (B) Quantitative analysis of seedling establishment in 14-day-old WT and *kam2-2* seedlings as shown in (A). Error bars show binomial 95% confidence intervals. Different letters indicate significant differences between three samples within the same genotypes at *p* < 0.01 (Fisher’s exact test with a Bonferroni multiple testing correction for each genotype). *n* = 134 (WT, +Suc), 133 (WT, –Suc), 139 (WT, +Sheet), 131 (*kam2-2*, +Suc), 133 (*kam2-2*, –Suc), and 140 (*kam2-2*, +Sheet).

### The primary cause of seedling arrest in membrane-trafficking mutants is not the lack of a carbon source

Previous studies have revealed sucrose-dependent seedling establishment in several membrane-trafficking mutants besides the *kam2*/*grv2* mutants (Kleine-Vehn et al., 2008; Thazar-Poulot et al., 2015; Zwiewka et al., 2011). We tested whether the use of plastic sheets would also rescue seedling arrest in these other mutants under sucrose-deficient conditions. We grew three membrane-trafficking mutants, *pat4-2*, *snx1-2*, and *mag1-1*/*vps29-1*, on sucrose-free or sucrose-containing medium (0.5% gellan gum) and on sucrose-free medium (0.5% gellan gum) covered with plastic sheets (Fig. 5A). The wild type showed >95% seedling establishment under all three conditions (Fig. 5B). By contrast, the membrane-trafficking mutants had a seedling establishment rate of more than 99% on sucrose-containing medium and a reduced seedling establishment rate on sucrose-free medium, which was recovered to some extent by the use of plastic sheets (Fig. 5B). These results suggest that carbon sources are not necessarily required for seedling establishment in these membrane-trafficking mutants and that seedling growth arrest is influenced by the growth environment after germination.

**Fig. 5.**
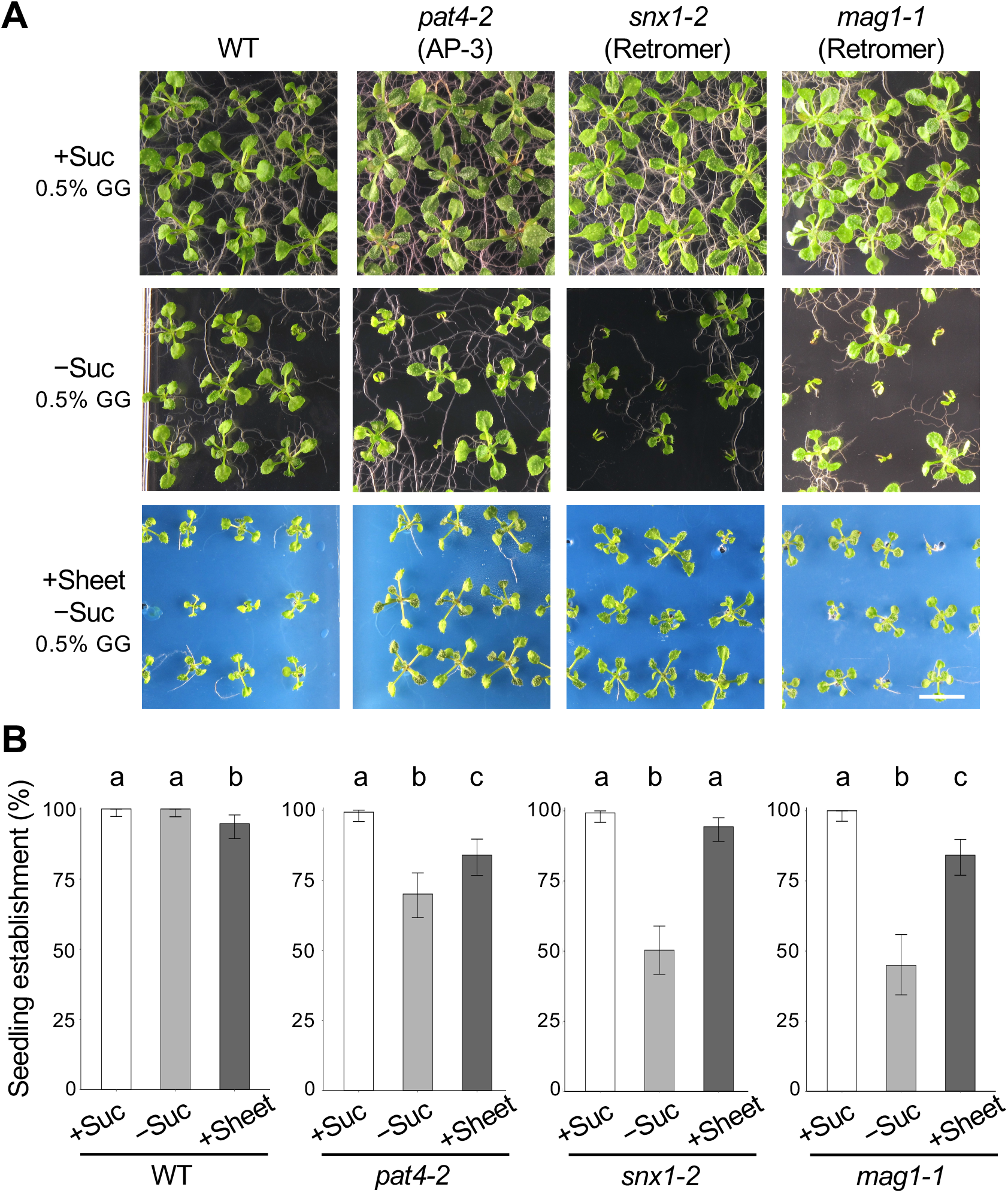
Applying a plastic sheet on solid medium restores the seedling establishment of other membrane-trafficking mutants. (A) Images of 14-day-old wild-type (WT), *pat4-2*, *snx1-2*, and *mag1-1*/*vps29-1* (*mag1-1*) seedlings grown on solid culture medium (0.5% gellan gum, GG) with 1% sucrose (+Suc), without sucrose (–Suc), or without sucrose and covered with a plastic sheet (+Sheet). Bar, 1 cm. (B) Quantitative analysis of seedling establishment in 14-day-old WT, *pat4-2*, *snx1-2*, and *mag1-1* seedlings as shown in (A). Error bars show binomial 95% confidence intervals. Different letters indicate significant differences between three samples within the same genotypes at *p* < 0.05 (Fisher’s exact test with a Bonferroni multiple testing correction for each genotype). *n* = 134 (WT, +Suc), 138 (WT, –Suc), 138 (WT, +Sheet), 135 (*pat4-2*, +Suc), 137 (*pat4-2*, –Suc), 137 (*pat4-2*, +Sheet), 134 (*snx1-2*, +Suc), 139 (*snx1-2*, –Suc), 141 (*snx1-2*, +Sheet), 97 (*mag1-1*, +Suc), 89 (*mag1-1*, –Suc), and 139 (*mag1-1*, +Sheet).

### RNA-seq analysis reveals changes in gene expression based on the rate of seedling establishment

To explore why the *kam2* mutant undergoes seedling arrest on sucrose-free medium at the transcription level, we examined global gene expression by RNA sequencing (RNA-seq) analysis. Three growth conditions were used: 0.5% gellan gum medium (normal seedling establishment for the wild-type seedlings), 0.2% gellan gum medium (reduced seedling establishment for both the wild-type and *kam2* seedlings), and 0.5% gellan gum medium covered with plastic sheets (restored seedling establishment for *kam2* seedlings), all on sucrose-free medium. To monitor gene expression variations as early as possible, we used seedlings at 4 days after sowing, when it was a little too early to determine with certainty whether they were undergoing growth arrest (Fig. 6A). We extracted RNA from a pool of 20 seedlings each of the wild type and *kam2-2* plants per condition, and five biological replicates were performed. In total, transcriptomes of 30 samples were acquired. Sufficient reads were obtained from each sample for analysis, with an average number of mapped reads of 10.5 million/sample (Supplementary Fig. S1, Supplementary Table S1). After normalization of the gene expression data (Supplementary Fig. S2), obvious differences in gene expression profiles between sample conditions were evident (Supplementary Fig. S3). Principal component analysis (PCA) revealed that individual samples of biological replicates clustered in close proximity to each other (Fig. 6B). The first and second principal components explained 29% and 19% of the variance, respectively. The first principal component separated groups of plants with different seedling establishment percentages under different growth conditions without plastic sheets, while the second principal component separated groups of plants under different growth conditions with and without plastic sheets. In particular, we observed similar transcriptomes in wild-type plants grown on 0.2% gellan gum medium and *kam2-2* plants grown on 0.5% gellan gum medium, both without plastic sheets (Fig. 6B), which showed similar seedling establishment rates (Fig. 2B).

**Fig. 6.**
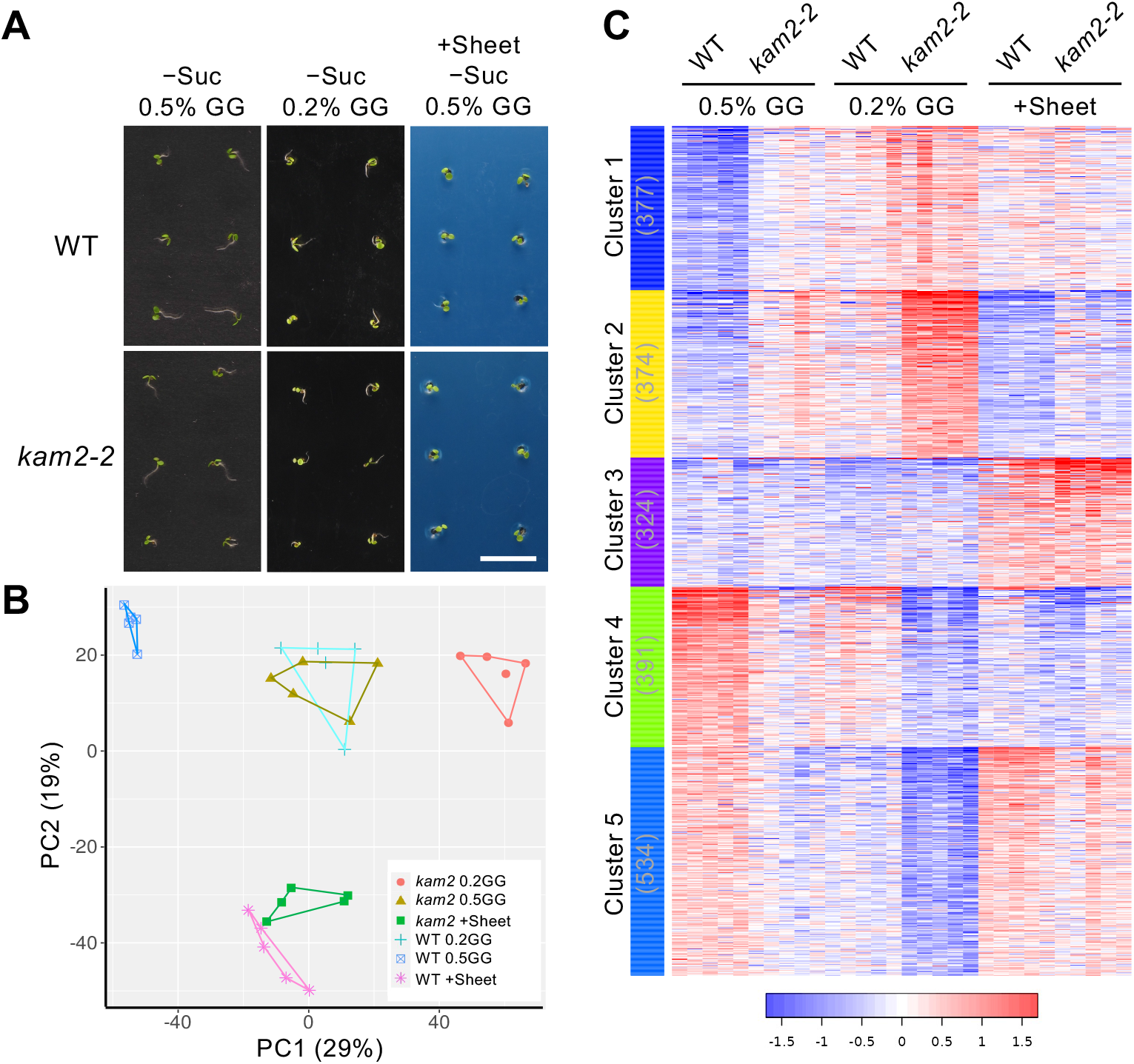
RNA-seq analysis of wild-type and *kam2* plants. (A) Images of 4-day-old wild-type (WT) and *kam2-2* seedlings grown on 0.5% gellan gum (GG) culture medium (0.5% GG), 0.2% GG culture medium (0.2% GG), and 0.5% GG culture medium covered with plastic sheets (+Sheet), all on sucrose-free medium. Total RNA extracted from these seedlings was used for RNA-seq. Bar, 1 cm. (B) Principal component analysis (PCA) of gene expression values of 4-day-old WT and *kam2-2* seedlings grown on solid culture medium as shown in (A). Symbols with different shapes and colors represent the three different growth conditions and genotypes as shown in the inset. The same symbols indicate the five biological replicates of the growth condition/genotype. Each symbol represents the expression value from 20 seedlings. (C) A heat map of k-means cluster analysis (k = 5) of the expression profiles of the top 2,000 variable genes across 30 samples (five replicates of six different samples). The horizontal axis indicates the growth conditions shown in (A), and the left vertical axis shows the five clusters, with the number of genes in parentheses. The color scale represents the relative expression levels of genes: red, blue, and white indicate high, low, and moderate relative expression levels, respectively.

To capture gene expression related to seedling arrest, we performed cluster analysis using the top 2,000 genes showing expression variation. K-means clustering classified the genes into five groups based on gene expression profiles (Fig. 6C, Supplementary Table S2). We performed Gene Ontology (GO) enrichment analysis of the groups of genes in each cluster (Fig. 7, Supplementary Table S3). Cluster 1 genes were expressed at low levels in the wild type on normal 0.5% gellan gum medium but at high levels in *kam2* on 0.2% gellan gum medium. These genes were enriched for the response to decreased oxygen levels (Supplementary Table S3A). Cluster 2 genes were expressed at low levels in the wild type on normal 0.5% gellan gum medium and at high levels in *kam2* on 0.2% gellan gum medium but were less highly expressed than cluster 1 on medium covered with plastic sheets. Cluster 2 was enriched for genes related to response to salicylic acid and plant defense response (Supplementary Table S3B). Cluster 3 gene expression increased in plants grown on medium covered with plastic sheets (Supplementary Table S3C). Cluster 4 genes were highly expressed in the wild type on normal 0.5% gellan gum medium but were downregulated in *kam2* on 0.2% gellan gum medium; these genes were enriched for S-glycoside (glucosinolate) biosynthetic process (Supplementary Table S3D). Cluster 5 genes were highly expressed in the wild type on normal 0.5% gellan gum medium but were downregulated in *kam2* on 0.2% gellan gum medium and were highly expressed on medium covered with plastic sheets. These genes were enriched in the GO terms response to auxin and cell wall organization (Supplementary Table S3E), which is consistent with the better seedling establishment under these growth conditions. Our results indicate that transcriptomic changes dependent on the rate of seedling establishment occurred before growth arrest could be detected.

**Fig. 7.**
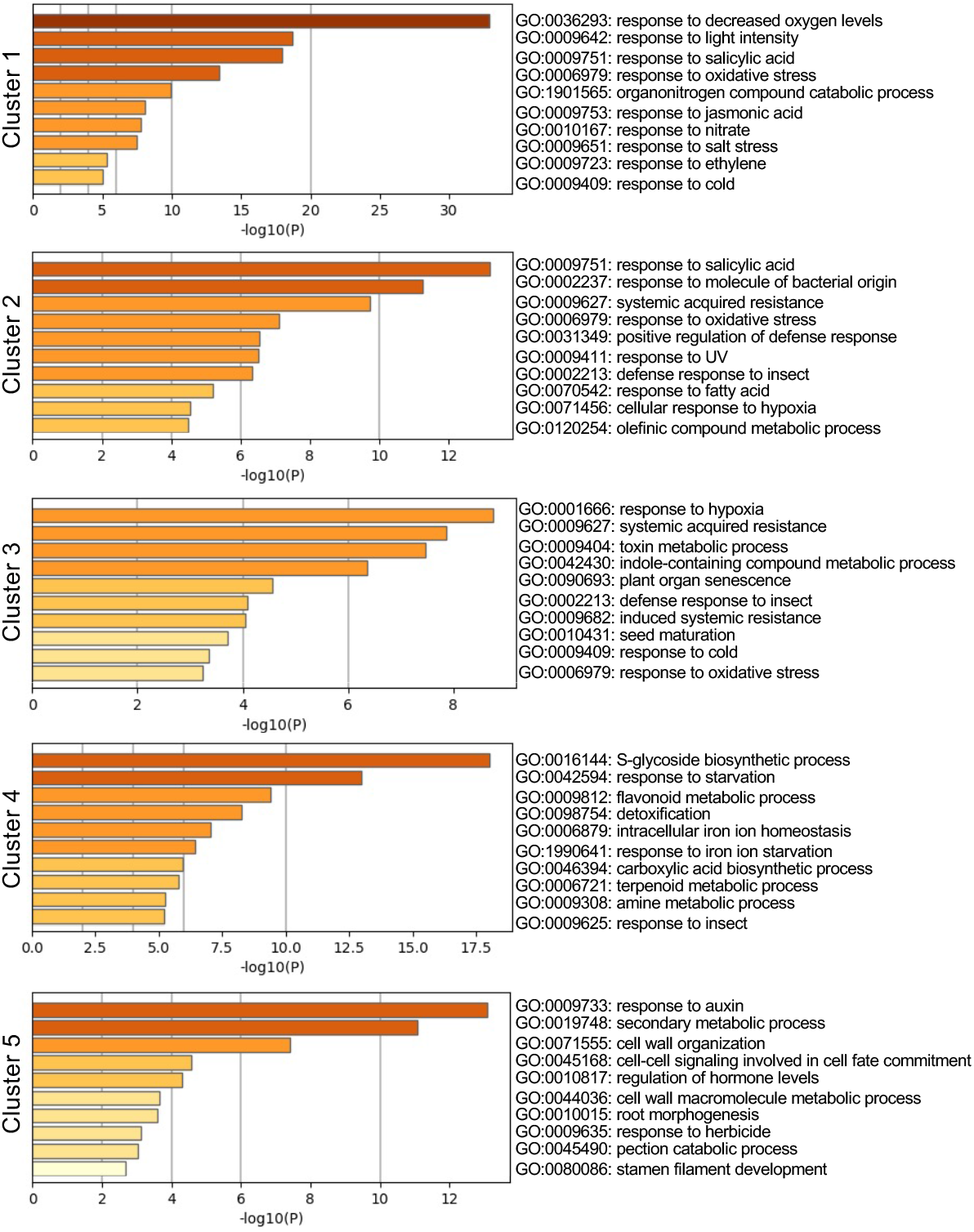
Gene Ontology (GO) enrichment analysis of the five clusters. GO analysis of biological processes was performed using genes in the five clusters shown in Figure 5C. The top 10 GO categories, color-coded by *p* values, are shown for each cluster. The horizontal axis indicates significance (–log10(*p*)).

Next, we examined the expression levels of genes identified in the above RNA-seq analysis in individual seedlings. Total RNA was extracted from 10 wild-type seedlings grown on 0.2% gellan gum medium without sucrose 4 days after sowing. The establishment rate of wild-type seedlings grown under this condition was around 45–66% (Figs. 2, 4), but it was unknown which individuals would stop growing. RT-PCR analysis showed that the expression level of the control *ACTIN2* gene was almost constant in all individuals but that the expression levels of genes in GO clusters 1 and 5 differed among individuals (Supplementary Fig. S4). The two genes in cluster 1 (AT3G47340 and AT4G35770) showed similar expression patterns within each seedling, while the expression of AT5G38910 in cluster 5 did not follow the same expression pattern as the two cluster 1 genes. These results suggest that individual seedlings differ in gene expression depending on subsequent seedling growth.

## Discussion

This study was prompted by the observation that a portion of mutants of the membrane-trafficking factor KAM2/GRV2 require sucrose for seedling establishment, as growth arrest occurs after germination on sucrose-free medium (Silady et al., 2008). As not all *kam2* seedlings show growth arrest in the absence of sucrose in the medium (Fig. 1A, B), we reasoned that the growth arrest is due to subtle differences in the growth environment (other than the lack of a carbon source), in addition to a genetic factor. Indeed, when we transplanted the arrested *kam2* mutants into fresh sucrose-free medium, the arrested seedlings resumed growth to some extent (Fig. 1D, E). Covering the sucrose-free medium with plastic sheets also restored seedling establishment (Fig. 3). As the plastic sheet prevents the shoots from contacting the medium (Fukuda et al., 2016), we speculated that the seedling arrest of *kam2* is due to contact with the medium. We observed a similar seedling arrest phenotype in wild-type plants grown on softer medium (Fig. 2). Since shoots are more likely to come in contact with the medium at lower gel concentrations (Fukuda et al., 2016), perhaps the growth arrest of wild-type seedlings also occurred due to contact with the medium. These results suggest that seedling arrest may be a universal stress response in Arabidopsis and is not specific to membrane-trafficking mutants. We observed poor seedling establishment in wild-type plants grown on 0.2% gellan gum medium and in *kam2-2* plants grown on 0.5% gellan gum medium (Fig. 2A, B). RNA-seq analysis showed that these plants had similar transcriptomes (Fig. 6B), strengthening the notion that the growth arrest exhibited by wild-type and *kam2* seedlings is the same stress response. It is currently unclear why *kam2* seedlings exhibit a hypersensitive growth arrest response compared with wild-type plants on 0.5% gellan gum sucrose-free medium. One possibility is that they are more likely to come in contact with the medium, and another possibility is that they are more likely to exhibit growth arrest when they come in contact with the medium.

The *kam2* mutant (and wild-type Arabidopsis) showed growth arrest at a stage when the cotyledons had expanded but the seedlings had not yet developed true leaves (Figs. 1 and 2). This is the stage when plants transition from heterotrophic growth, using seed storage materials, to autotrophic growth, relying on photosynthesis by their expanded cotyledons (Ha et al., 2017). Therefore, the *kam2* mutant exhibits defects in this phase transition. When the cotyledons are unable to acquire sufficient photosynthetic capacity, seedlings may temporarily stop growing until the environment improves as an adaptive stress response. The transition to autotrophic growth is a key turning point in plant development; there may be a checkpoint controlling this phase transition. Similar to *kam2*, several other Arabidopsis mutants defective in this phase transition have been reported: *à bout de souffle* (Lawand et al., 2002), *stimpy* (Wu et al., 2005), *wall-associated kinase2* (Kohorn et al., 2006), *wrinkled1* (Cernac et al., 2006), *calmodulin like protein39* (Bender et al., 2013), *gassho1 gassho2* (Racolta et al., 2014), *ndufs4* (Kühn et al., 2015), and *kerberos* (Doll et al., 2020). These mutants do not form true leaves and arrest growth when germinated on sucrose-free medium but form true leaves and grow normally on sugar-containing medium. The causal genes of these mutants encode transcription factors, signaling factors, or mitochondrial proteins. However, the specific functions of these genes in seedling establishment are not well understood. As with *kam2*, the frequency of the growth arrest phenotype of these mutants on sucrose-free medium is not 100%, suggesting that both environmental and genetic factors play a role. It would be interesting to determine whether changing the concentration of gel in the sucrose-free medium, applying plastic sheets to the sucrose-free medium, or performing transplantation experiments from a sucrose-free medium to a fresh sucrose-free medium would alter the rate of seedling establishment in these mutants.

Although we determined that shoot contact with the medium causes the growth arrest of *kam2* and wild-type Arabidopsis seedlings, the underlying physiological and genetic mechanisms are currently unclear. What happens in the plant body after shoots touch the medium? RNA-seq data could provide clues about the transcriptional status of growth-arrested plants (Figs. 6, 7). GO enrichment analysis showed that cluster 2 was enriched in genes for salicylic acid and defense-related responses, while cluster 5 contained many growth-related genes, such as genes related to the response to auxin and the cell wall (Fig. 7). Since there is a trade-off between plant growth and defense responses (Margalha et al., 2019), the growth arrest in *kam2* seedlings may result from the induction of defense responses. However, we cannot rule out the possibility that growth arrest and defense responses occur independently in the mutant. The arrested *kam2* mutant may be under hypoxia stress (Fig. 7). Hypoxic responses are often observed in submerged plants, which trigger defenses responses; there is an elevated risk of infection by pathogens after flooding (Hsu and Shih, 2013). The GO terms ‘response to decreased oxygen levels’ enriched in cluster 1 and ‘cellular response to hypoxia’ enriched in cluster 2 may be related to the seedling arrest phenotype. When plants are submerged due to flooding, photosynthesis is inhibited due to impaired CO_2_ absorption, and respiration is inhibited due to low O_2_ levels (hypoxia), making the plant unable to break down stored substances to produce energy (Cho et al., 2021). Some wetland plants subjected to flooding survive by pausing growth to conserve energy and resume development when environmental conditions are suitable (León et al., 2021). Seedlings cultured in soft solid medium tend to contain more water in the apoplast and are more likely to enter a state of hyperhydricity, which can induce hypoxia responses (van den Dries et al., 2013). Indeed, the arrested wild-type and *kam2* seedlings had translucent cotyledons and might be in a state of hyperhydricity (Figs. 1, 2). The Arabidopsis *atp-binding cassette transporter subfamily g5* (*abcg5*) mutant fails to develop true leaves and shows arrested seedling growth under waterlogged conditions (Lee et al., 2021). ABCG5 is involved in cuticle formation, which blocks the penetration of water into the plant body; *abcg5* plants show hyperhydricity and a hypoxia response.

The growth-arrested phenotype of the membrane-trafficking mutants *pat4* (Zwiewka et al., 2011), *snx1* (Kleine-Vehn et al., 2008), and *mag1*/*vps29* (Thazar-Poulot et al., 2015) were partially rescued when grown on plastic sheets on growth medium, as did *kam2* (Fig. 5). These results suggest that these membrane-trafficking mutants also exhibit seedling arrest in an unfavorable growth environment. PAT4 is a subunit of the Golgi-localized adaptor protein complex 3 (Zwiewka et al., 2011), while SNX1 and VPS29 are subunits of the retromer complex (Kleine-Vehn et al., 2008; Shimada et al., 2006). The seedling arrest commonly observed in these membrane-trafficking mutants is likely caused by abnormal post-Golgi transport. One candidate post-Golgi transport cargo protein is the auxin transporter PIN. These mutants also exhibit abnormal PIN protein transport and abnormal auxin responses (Jaillais et al., 2007; Kleine-Vehn et al., 2008; Zwiewka et al., 2011). The *grv2* mutant, an allelic mutant of *kam2*, also shows abnormal auxin-related gravitropism (Silady et al., 2004). Indeed, the GO term ‘response to auxin’ was enriched in genes in cluster 5, whose expression decreased when the rate of seedling establishment was reduced (Fig. 7). Many laboratories (including ours) use sucrose-containing solid medium to improve Arabidopsis germination and growth. Perhaps various mutants (including membrane-trafficking mutants) should be grown on sucrose-free or plastic sheet-covered medium to determine whether they are defective in phase transitions. Growth arrest, which was previously thought to be caused by a shortage of lipids or metabolic abnormalities, might be an environmental response; we hope that future research will advance our understanding of this phenomenon.

## Materials and Methods

### Plant materials and growth conditions

The *Arabidopsis thaliana* accession Columbia-0 (Col-0) was used as the wild type throughout the study. Details about the T-DNA insertion mutants *kam2-2* and *kam2-4* were previously reported (Tamura et al., 2007). The *mag1-1*/*vps29-1* mutant was previously reported (Shimada et al., 2006). *pat4-2* (SALK_069881) and *snx1-2* (SALK_033351) seeds were obtained from the Arabidopsis Biological Resource Center. The Arabidopsis seeds were surface-sterilized with 70% ethanol, sown on solid medium, and germinated after a 2-day incubation at 4°C in the dark to break dormancy. All plants were grown on half-strength Murashige and Skoog medium containing 1.0% (w/v) sucrose or without sucrose, 0.05% (w/v) MES-KOH (pH 5.7), and 0.2%, 0.5%, or 1% (w/v) gellan gum (FUJIFILM Wako) in a growth chamber under continuous white fluorescent light at 22°C. To quantify seedling establishment, 70 seeds were sown per dish using a plastic rectangular dish (140 × 100 × 14.5 mm, AW2000, EIKEN CHEMICAL). For experiments in which the solid medium was covered with a plastic sheet, a thin plastic file folder (0.2-mm thickness) with 1-mm-diameter perforations (50 holes per sheet) was placed on the surface of sucrose-free medium, and the seeds were sown in the hole. Seedling establishment was defined as the successful growth of seedlings with developing true leaves and elongated roots 14 days after incubation. The regrowth rate after transplantation was defined as the proportion of seedlings that had grown roots and rooted into the medium by 10 days after transplantation. Since the first and second true leaves of many of the individuals were somewhat expanded after transplanting onto sucrose-free medium, we focused on root elongation rather than leaf development to evaluate regrowth.

### DNA extraction from seedlings

Seedlings were homogenized with 3-mm plastic beads (YTZ-3, AS ONE) in DNA extraction buffer (200 mM Tris-HCl pH 7.5, 250 mM NaCl, 25 mM EDTA, and 0.5% [w/v] SDS). The extract was centrifuged at 20,000 g for 5 min at room temperature to remove the precipitate. An equal amount of isopropanol was added to the supernatant, and the sample was incubated for 2 min and centrifuged at 20,000 g for 5 min at room temperature. After removing the supernatant, 70% EtOH was added to the precipitate, and the sample was centrifuged at 20,000 g for 5 min at room temperature. After the supernatant was removed, the precipitate was dried and dissolved in 30 µL of TE buffer (10 mM Tris-HCl pH 8.0 and 1 mM EDTA).

### Genotyping of T-DNA insertion mutants

The genotyping of T-DNA insertion mutants was performed by PCR with genomic DNA and gene- and T-DNA-specific primers. The sequences of the primers were as follows: for *PAT4*, gene-specific primers 5□-GTCTTAACGCCTTGTCGATTG-3□ and 5□-TGCAGTTGCAGAAATAGGACC-3□; for *SNX1*, gene-specific primers 5□-TCAAGCACCCAAAAGCATTAC-3□ and 5□-TGGACAGATTCAGGTTTCAGG-3□; for the T-DNA-specific primer LBb1.3 5□-ATTTTGCCGATTTCGGAAC-3□. PCR was performed at 95°C for 2 min, followed by 35 cycles at 98°C for 10 sec, 55°C for 15 sec, and 68°C for 30 sec, using the PCR enzyme KOD FX Neo (TOYOBO).

### RNA-seq and GO enrichment analyses

Total RNA was extracted from wild-type and *kam2* seedlings grown on solid medium (0.2 or 0.5% gellan gum) or solid medium (0.5% gellan gum) covered with plastic sheets, all on sucrose-free medium at 4 days after sowing. The seedlings were snap-frozen in liquid nitrogen and homogenized with 6-mm plastic beads twice at 500 strokes/sec for 20 sec. After adding 500 µL of 1-thioglycerol/homogenization solution on ice, the sample was incubated at room temperature for 2 min in a shaker. The samples were centrifuged at 14,000 g for 2 min, and supernatants were aspirated into 1.5-mL tubes containing lysis buffer. The samples were vortexed for 15 sec, incubated at room temperature for 10 min, and centrifuged at 14,000 g for 4 min. RNA was extracted from pools of 20 seedlings of each genotype grown under the three conditions, with 5 replicates each using a Maxwell RSC Instrument. An RNA-seq library was prepared using the Lasy-Seq v1.1 protocol (Kamitani et al., 2019). The library was sequenced using the 150-bp paired-end read mode on the HiSeqX platform (Illumina, CA, USA). Sequenced single-end reads were preprocessed and mapped to the Arabidopsis genome (TAIR10) using the web tool RaNA-seq v.1.0 (Prieto and Barrios 2020, https://ranaseq.eu/). The mapped data were quantified, and PCA and k-means cluster analyses were performed using the web tool iDEP v.0.96 with default settings unless otherwise specified. (Ge et al., 2018) (http://bioinformatics.sdstate.edu/idep/). GO analysis was performed with Metascape v.3.5.20230501 (Zhou et al., 2019) (https://metascape.org/gp/index.html#/main/step1).

### RT-PCR analysis

Total RNA was isolated from 4-day-old wild-type seedlings using a Maxwell RSC Plant Kit (Promega). cDNA was synthesized from total RNA (100 ng) using ReverTra Ace qPCR RT Master Mix with gDNA Remover (TOYOBO). PCR was performed using the cDNA and KOD One Master Mix (TOYOBO). PCR conditions were as follows: 98°C for 1 min, 30∼40 cycles of 98°C for 10 sec, 60°C for 5 sec, and 68°C for 5 sec. The PCR products were separated by 1.0% (w/v) agarose gel electrophoresis and stained with Midori Green Extra (NIPPON Genetics). The DNA bands were detected with LAS-Digi PRO (NIPPON Genetics). The primers used for amplification of each gene are as follows: 5□-CAAGCCAGAGGAAGGAAGGATC-3□ and 5□-TATTGTTGGACCAGCTTGCATC-3□ (for AT3G47340), 5□-GCCAATTACCAACACGAAGAGG-3□ and 5□-GTTTCCTGAAGTGGGACGAGAC-3□ (for AT4G35770), 5□-AGCTCTCTTCAGGACTTTTGCG-3□ and 5□-AGACATCACCCTCGTAAAGTGT-3□ (for AT5G38910), and 5□-AGAGATTCAGATGCCCAGAAGTCTTGTTCC-3□ and 5□-AACGATTCCTGGACCTGCCTCATCATACTC-3□ (for *ACT2*).

### Statistical analysis

Statistical analyses were performed in R version 4.2.1. Fisher’s exact tests in the package “multicomp” were used for multiple comparisons of different growth conditions and genotypes.

## Supplementary Data

Supplementary data are available at PCP online.

## Data Availability

The RNA-seq data are available at the DDBJ Sequence Read Archive (DRA) under the accession number DRA016745.

## Disclosure

The authors have no conflicts of interest to declare.

