## Supplementary material for "The Arabidopsis *katamari2* Mutant Exhibits a Hypersensitive Seedling Arrest Response at the Phase Transition from Heterotrophic to Autotrophic Growth": Fig. S1

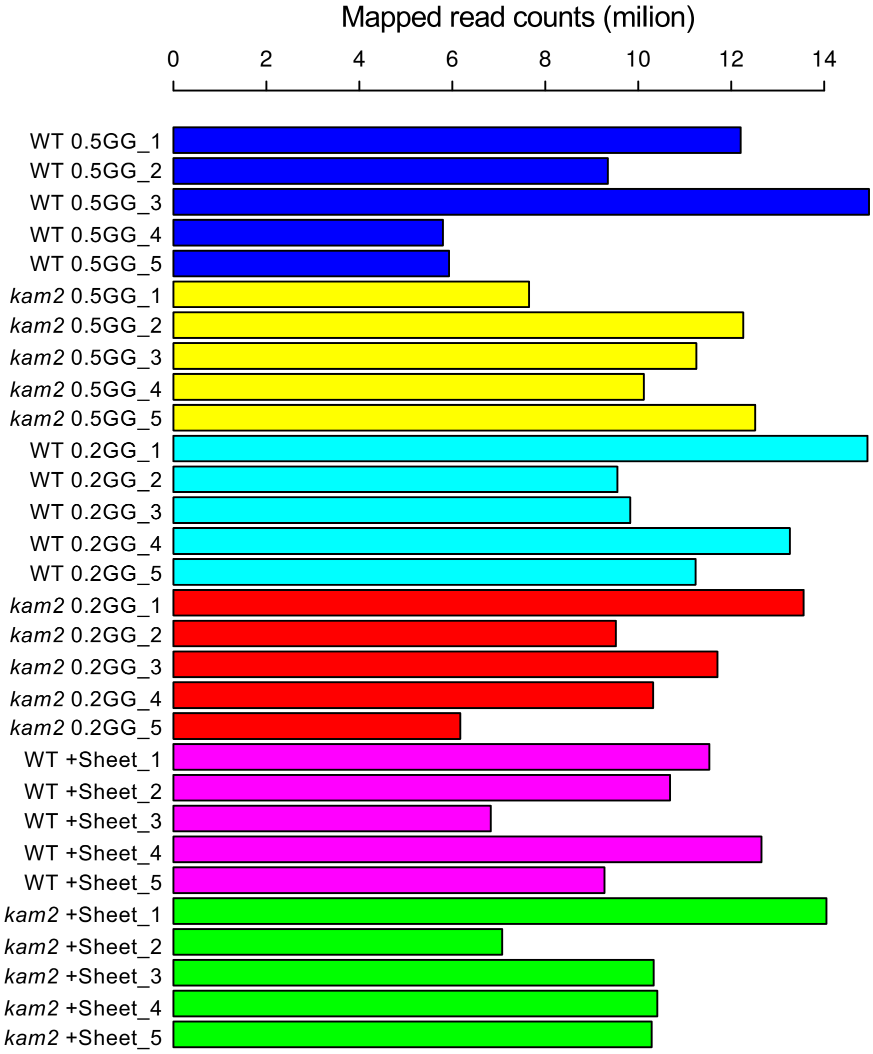


**Supplementary Fig. S1** Total mapped-read counts of 30 samples (five replicates of six different samples): wild-type (WT) and *kam2-2* seedlings grown on 0.5% gellan gum (GG) medium (0.5GG), 0.2% GG medium (0.2GG), and 0.5% GG medium covered with plastic sheets (+Sheet), all on sucrose-free medium.
