## Supplementary material for "The Arabidopsis *katamari2* Mutant Exhibits a Hypersensitive Seedling Arrest Response at the Phase Transition from Heterotrophic to Autotrophic Growth": Fig. S4

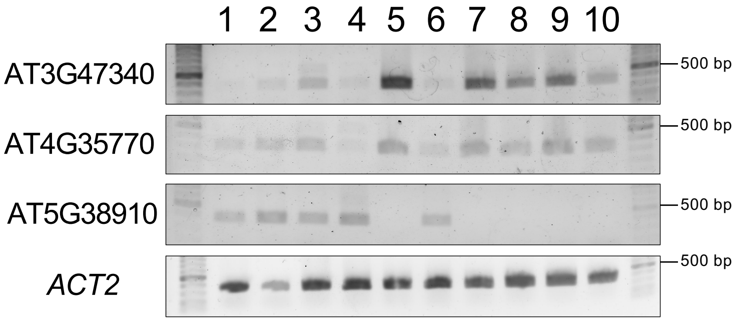


**Supplementary Fig. S4** RT-PCR analysis showing differential gene expression in each seedling depending on subsequent seedling growth. Electrophoresis images of RT-PCR products of 4-day-old wild-type seedlings grown on solid culture medium containing 0.2% gellan gum without sucrose. Total RNA from 10 individual seedlings was subjected to reverse transcription. *ACTIN2* (*ACT2*) was used as an internal control. The number of PCR cycles used to amplify each gene was as follows: 30 (AT3G47340, AT4G35770, *ACT2*) and 40 (AT5G38910).
